# Freshwater degradation as a potential extinction factor of the Japanese river otters

**DOI:** 10.1101/2025.10.30.685517

**Authors:** Kanzi M. Tomita, Tamihisa Ohta, Phillip J. Manlick, Shingo Kobayashi, Chikage Yoshimizu, Ichiro Tayasu

**Affiliations:** Faculty of Agriculture and Marine Science Kochi University, Kochi, Japan; Faculty of Science, Academic Assembly, University of Toyama, Toyama, Japan; Pacific Northwest Research Station, USDA Forest Service, Juneau, AK, USA; Ehime Prefectural Science Museum, Ehime, Japan; Research Institute of Humanity and Nature, Kyoto, Japan

**Keywords:** sulfur isotopes, species extinction, Mustelidae, museum specimen, freshwater conservation

## Abstract

Exploring the causes of species extirpation can provide insights into mitigating the ongoing extinction. The Japanese otter *Lutra lutra* whiteleyi had extincted around 1970s, but their extinction causes remain unknown. To elucidate the ecological relationship among their foraging ecology, habitat degradation, and the extinction, we assessed dependency of Japanese otters on marine- or freshwater-derived resources using sulfur stable isotopes (*δ*^34^S: ^34^S/^32^S) of the museum specimens which were collected on the brink of extinction. Our results clearly showed that the otters strongly depended on marine-derived resources, indicating anthropogenic activities led to depletion of freshwater prey and a potential extinction factor of the last population of otters in Japan. Protecting freshwater habitats was crucial for preventing otter extinction in Japan, and for safeguarding the extant otter species.

## Introduction

Many wild animals have threatened extirpation due to anthropogenic pressures such as biological invasion, overexploitation and land use intensification (Ceballos et al. 2015, Schmitt et al. 2018, Wiens and Saban 2025). Exploring the causes for the extirpation can provide insights into mitigating the ongoing extinction because the past extinctions is likely to share the same drivers with contemporary species loss (Novacek and Cleland 2001, Andermann et al. 2020, Smith 2021, Mychajliw et al. 2024). However, it can be challenging to reconstruct ecological information such as diet, movement or trophic niche for organisms that have been extirpated.

Ecosystem connectivity and intactness are essential for animals that utilize multiple ecosystems (Rapport et al. 1998, Carranza et al. 2012, Doherty and Driscoll 2018, Brodie et al. 2025). Loss of ecosystem connectivity and intactness are critical for resting, breeding, and foraging activities of these animals and can lead to species decline or extirpation by forcing them to rely on suboptimal or alternative ecosystems (Beger et al. 2010, Heinrichs et al. 2016). Local extirpation due to anthropogenic environmental changes have occurred both in the past and the present worldwide (Turvey and Crees 2019, Andermann et al. 2020). Thus, elucidating the relationships among ecosystem degradation, ecological responses of animals and the past extirpation can help to predict risk of future extinction (Campbell et al. 2022).

The otter (Mustelidae; Lutrinae) is a semiaquatic mammal of the mustelid family There are 13 known species in the world, all of which are endangered (Kruuk 2006). Almost species, except for sea otters (*Enhydra lutris*) and sea cats (*Lontra felina*), depend on both terrestrial and aquatic habitats as nesting and foraging sites. Seven species utilize both freshwater and marine ecosystems and its preference differs among species and regions (Beja 1991, Clavero et al. 2006, Kruuk 2006). Otters have been threatened by anthropogenic disturbances such as land development and contamination (Roos et al. 2001, Jo et al. 2017). In particular, pollution of freshwater habitats such as emission of chemical compounds and heavy metals can not only impair otter health but also decline food availability (Sjöåsen et al. 1997, Roos et al. 2001, Klenavic et al. 2008). Therefore, otter conservation requires maintaining the integrity of multiple interconnected ecosystems across land, marine, and river.

In Japan, a subspecies of Eurasian otter (*Lutra lutra* whiteleyi) had extincted around 1970s (Ando 2008, Naito et al. 2015, Waku et al. 2016). The causes of extinction are commercial fur hunting, poaching, net entanglement in coastal fishery, and water pollution (Ando 2008). From the late 19 to early 20 centuries, Japanese otters were extensively hunted due to increased fur demand during wartime. Nevertheless, freshwater pollution is closely associated with otter extinction in Japan because water pollution by industrial wastewater and pesticides (e.g., PCBs and DDTs), river improvement, and dam construction were intensive during the high economic growth period (the mid 20 century), coincided with otter extinction in Japan (Itsukushima et al. 2021). In this period, many freshwater fishes were registered as endangered species and some species had extincted locally throughout Japan (Nakagawa and Mori 2025). River fragmentation by dam construction can markedly decrease freshwater fish abundance (Fukushima et al. 2007). Therefore, elucidating whether Japanese otters relied primarily on freshwater or marine food resources during the terminal phase of their population is crucial for understanding the drivers of their extinction and informing future conservation strategies for extant otter species.

We predicted that Japanese otters on the brink of extinction substantially depended on marine derived resources due to degradation of freshwater ecosystem. We assessed dependency of Japanese otters on marine-derived resources using sulfur stable isotopes (*δ*^34^S: ^34^S/^32^S) of the museum specimens which were collected on the brink of extinction. The *δ*^34^S value is an indicator for estimating whether river- or marine-derived resources significantly affect an organism’s body composition (Fry and Chumchal 2011, MacAvoy et al. 2015, Raoult et al. 2024, Tanaka et al. 2025). It represents high value in marine environments (Rees et al. 1978), whereas it shows much lower values in freshwater ecosystems (Fry and Chumchal 2011). Our primary goal is to elucidate which ecosystem Japanese otters relied on food resources based on *δ*^34^S values of the museum specimens. We further explored ecological factors affecting *δ*^34^S values by building multiple regression models combining with sex, capture year, and freshwater accessibility of the specimens.

## Methods

### Study site

This study were conducted on the southwestern part of the Shikoku Island, southern Japan (Fig.1a) where is one of the last confirmed sightings of river otters in Japan. Otters had been observed in the mid- and upper parts of rivers in the study area until around 1930s (Ando 2008). The representative riverine system in the study site had degraded due to river improvement and intensive use of pesticides (i.e., DDTs and PCBs) for cultivation of mandarin oranges and rice since 1940s. Freshwater fauna in the study site would deplete in this period, given that they declined by these anthropogenic disturbances throughout Japan (Nakagawa and Mori 2025). The studied population in southwestern Shikoku Island had been rediscovered along shoreline in 1950s. This area is characterized by a typical ria coastline with highly complex shorelines and multiple inner bays (Figure1a), suggesting that such topographic features likely fulfilled the habitat requirements of otters (Kruuk 2006). Otters had been preserved legally by the local government since 1960s when they were mostly extincted. However, otters have not been observed in this area since 1975, and this is the last local population of otters throughout Japan.

**Figure 1.**
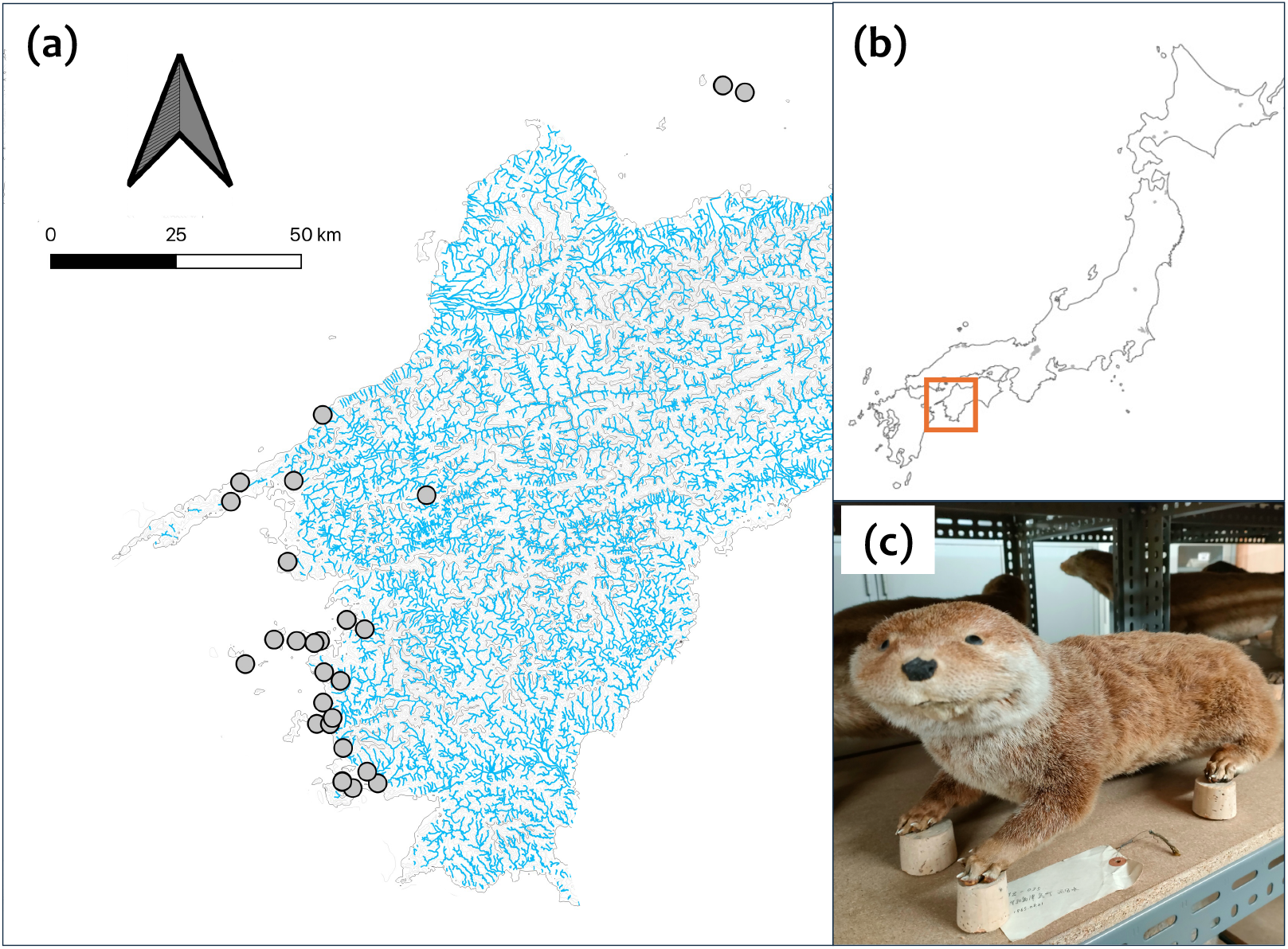
**(a)** Sampling points of otter *<u>Lutra lutra whiteleyi</u>* specimens in the southwestern part of Shikoku Island. Blue bold lines indicate riverine areas based on data from the National Land Numerical Information (https://nlftp.mlit.go.jp/ksj/gml/datalist/KsjTmplt-W05.html). **(b)** Study site location on Japan (orange square) **(c)** A sampled *L. lutra whiteleyi* specimen displayed in the Ehime Prefectural Science Museum.

### Specimen sampling

We sampled body hair (almost guard hair) and vibrissae of a total of 36 otter specimens (32 taxidermy and 2 fur) for isotopic analysis from the collection of the Ehime Prefectural Science Museum (Ehime, Japan) in April 2024 (Figure1a). These specimens were collected from 1954 to 1975, on the brink of extinction of otters in Japan. Four individuals were captured on islands (Figure 1b). Specimens were identified as *L. lutra* whiteleyi from characteristics of rhinarium and tail-body ratio (Ando 2008). Thirty-one specimens were made from otters which were collected by a conservation project led by Ehime prefecture. The remaining 5 specimens were donated from local public schools and failed to information on date, year, sampling locations, and sex. Mortality factors of otter specimens were recorded 27 cases: entanglement by fishery nets (N=20, 52.6 %), trapping for conservation (N=4, 10.5%), and natural death (N=3, 7.9%). Average body length of the specimens are 109.52 cm (SD = 12.89) and sex ratio is roughly equal (female=14, male =16, unknown =6). Based on the diet of otters in Japan (Ando 2008), we collected fishes and shells, and crabs as the potential prey items from freshwater (N=4) and marine (N=4) ecosystems around the study sites (Table S1). Muscles, soft bodies, and bulk bodies (exoskeleton, soft bodies, and innards) were sampled fish, shells, and crabs, respectively. We did not consider trophic levels of prey items because trophic discrimination can be ignored for sulfur stable isotopes (Raoult et al. 2024). All samples were soaked in deionized water for 24 h and dried at 60 C for >24 h and powdered by pestles and mortars.

### Sulfur stable isotope ra4os

Approximately 0.5 mg of otter samples, comprised of 5-10 hairs with 5-10 mm length of vibrissae from the base, were encapsulated in tin capsules, together with 1.0 mg of V_2_O_5_. In the case of food materials, approximately 2.5 mg with 5.0 mg of V_2_O_5_ were encapsulated in tin capsules. The weight of otter and food samples was determined based on the previous studies on sulfur isotope ratios of aquatic animals and terrestrial mammals (Funakawa et al. 2025, Tanaka et al. 2025). The stable isotope ratios of sulfur were measured using an EA-IRMS (EA Flash 2000, ConFlo IV, and Delta V Advantage, Thermo Fisher Scientific) at the Research Institute for Humanity and Nature (RIHN). The isotope ratios of sulfur are expressed in *δ* notation in accordance with the international standard scale (Vienna Cañon Diablo troilite) based on the following equation.

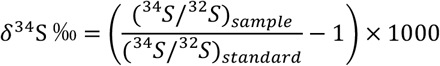

The *δ*^34^S data were calibrated against two international standards [IAEA-S-01 (*δ*^34^S= 0.3 ‰) and IAEA-S-02 (*δ*^34^S= 22.62 ‰)]. The analytical precision (± SD) for *δ*^34^S was calculated by the repeated measurements of working standards (per 8 samples), resulting in ± 0.12_‰_ in IAEA S-1 (n = 16) and ± 0.25_‰_ in IAEA S-2 (n = 18).

### Data analysis

We used *δ*^34^S values of vibrissae for the statistical analysis because of a strong positive correlation between otter hairs and vibrissae (r = 0.904, p <0.001). To investigate diet of the otters, we performed Bayesian stable isotope mixing models using the R package MixSIAR (Stock et al. 2018). We then compared the 95% highest probability density intervals among species. MixSIAR were set as follows: chain length = 4000,000, burn-in = 2000,000, thinning = 500 and three Markov chain Monte-Carlo methods. The discrimination value was set at −0.6 ± 1.3‰ (mean ± SD) based on a meta-analysis of *δ*^34^S of mammals (Raoult et al. 2024). The average *δ*^34^S values of marine and freshwater food items were 20.1 ± 1.43‰ and −2.7 ± 0.59 ‰, respectively (Table S1). Model convergence was assessed using the Gelman-Rubin and Geweke diagnostics. We considered model convergence when no variables had a Gelman-Rubin diagnostic of >1.05, and less than 5% of the variables were outside the 95% CI based on the Geweke diagnostic.

Since most specimens were collected along the coastline, freshwater accessibility may influence *δ*^34^S values. We created buffers around the sampling points and calculated the total river length contained within each buffer. Given movement ranges of otters (Kruuk 2006), we applied buffer sizes of 5, 10 and 20 km around the sampling points. These buffers do not mirror the availability of the accessible freshwater habitats because otters usually move along rivers and the amount of river discharge is not considered. However, the calculated values can be regarded as relative indexes of freshwater accessibility. We used river data from the National Land Numerical Information (https://nlftp.mlit.go.jp/ksj/gml/datalist/KsjTmplt-W05.html). We performed this spatial analysis using QGIS ver. 3.36 (QGIS Development Team 2024).

We constructed generalized linear models (GLMs) with a Gaussian error distribution to examine the effects of otter sex, capture year, and freshwater accessibility on *δ*^34^S values of otter vibrissae. We confirmed that the values fit a normal distribution according to the Kolmogorov-Smirnov test (p = 0.334). We constructed GLMs separately among 5, 10, and 20 km buffers. We did not use the average marine-derived resources estimated by Bayesian stable isotope mixing model because it was calculated a population level diet, but not individual-based diets. The explanatory variables were otter sexes, captured year, total river length within the buffers (freshwater accessibility), interaction terms between sex or captured year and river length. Along with our a priori hypothesis, capture year may positively influence *δ*^34^S values due to increased utilization of marine-derived resources. Prior to model fitting, we confirmed the absence of multicollinearity among explanatory variables by calculating the variance inflation factor (VIF). Model selection was conducted using the stepwise selection approach based on the Akaike Information Criterion (AIC). The final model was selected according to the lowest AIC value. Finally, to ensure the absence of the spatial bias in *δ*^34^S values, spatial autocorrelation was assessed using Moran’s I. All statistical analyses were conducted in R version 4.3.2 (R Core Team 2023), using the “stats”, “MASS”, “sf”, and “spdep” packages.

## Results

The average *δ*^34^S values of otter vibrissae were 17.3‰ (N=36, SD=1.11). Our Bayesian stable isotope mixing model was converged based on Gelman-Rubin diagnostic and Geweke diagnostic. The average marine- and freshwater-derived resources were estimated as 90.0% (95% CI: 0.831– 0.956) and 10.0% (95% CI: 0.044–0.169), respectively (Figure2), strongly indicating that the otters heavily depended on marine prey. For all buffer sizes, GLMs indicated no significant effects of otter sex and capture year on *δ*^34^S values. For 10 and 20 km buffer models, none of the explanatory variables showed a significant effect on *δ*^34^S value in the best models (Table S3 and S4). However, the best model with 5 km buffers suggested a marginally significant negative effect of total river length (Estimate = −0.007, SE = 0.004, *P*= 0.073) and the interaction between capture year and river length (Estimate = 3.57×10^−6^, SE = 1.90×10^−6^, *P* = 0.072) (Table.S2 and S5). This means that the *δ*^34^S value (otter’s dependence on marine-derived resources) decreased as freshwater accessibility around otter habitats increases. There was no spatial autocorrelation of *δ*^34^S values in the study sites (Moran’s I = 0.092, p = 0.156).

**Figure 2.**
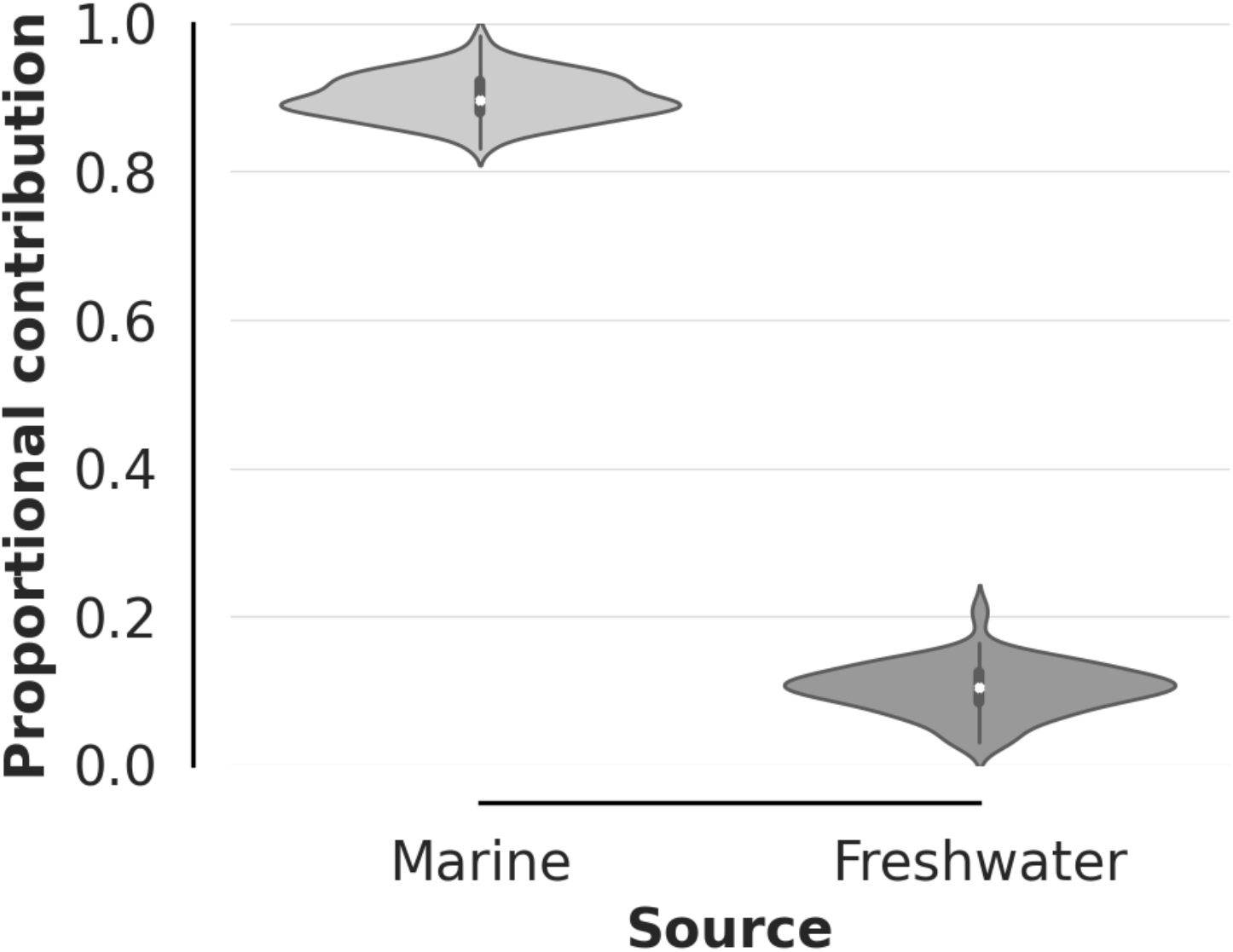
Violin plots showing the estimated proportional contributions of Marine and Freshwater resources based on *δ*^34^S values of vibrissae in the Japanese otters (N=36). The width of each violin represents the distribution density of posterior estimates, and the internal box plot indicates the interquartile range and median. Marine resources contributed substantially more than Freshwater resources.

## Discussion

Our results demonstrated that Japanese river otters on the blink of extinction depended mostly on marine-derived resources regardless of sex and body size, implying a link between otter extinction between their resource use. Furthermore, the significant interaction between river length and capture year suggests that the relationship between freshwater accessibility and reliance on marine-derived resources changed as the population neared extirpation (Figure 3). This result conform timing of declining freshwater preys due to anthropogenic degradation. There are three possible reasons for a highly dependence on marine-derived resources for the studied otters because almost specimens were captured around coastline; 1) inland population had extincted prior to coastline population; 2) inland population change their core habitats to coastline; 3) otters in the study area had originally lived along the coastline long before their extinction.

**Figure 3.**
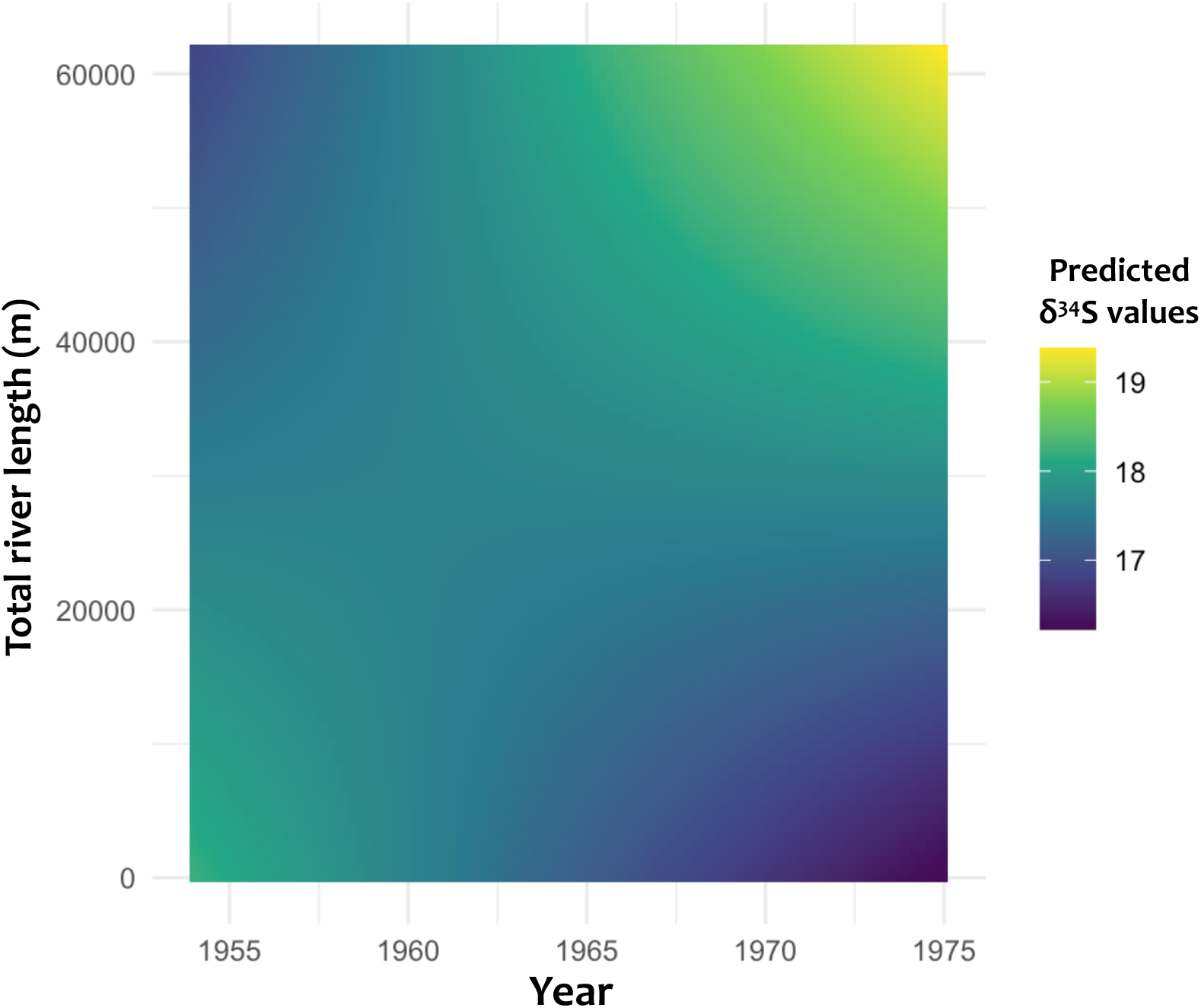
The heat map shows predicted values for sulfur stable isotopes (*δ*^34^S) of the Japanese otter *Lutra lutra* whiteleyi vibrissae based on the generalized linear model with interactive effects between total river length (meter) contained within 5km buffer around the sampling points and the captured year on *δ*^34^S values.

Inland areas were rapidly degraded due to human activities more severe than coastline areas in the study site during the high economic growth period. As freshwater degradation decrease prey and resting site availabilities for otters along rivers, otters might locally extirpated inland areas, or immigrated to coastline areas. The prior extirpation of inland population was also suggested by local records that otters were commonly observed at mid- or upper-parts of rivers in the study area before the study period (since 1950s) (Ando 2008). Japanese otters are supposed to utilize from freshwater to marine ecosystems, but primarily feed on freshwater foods such as fishes, snails, and crabs during the period when the population was stable (Ando 2008). Thus, freshwater degradation would change otter’s core habitats to coastline areas.

Because almost otter specimens were caught along coastline, they might have originally foraged along sea before near extinction in southwestern Shikoku Island as well as Eurasian otters in Shetland (Kruuk and Moorhouse 1990). However, in the study area, the coastline was unlikely to constitute the core habitats of the Japanese otter. The otters had been commonly observed in the mid- and upper parts of rivers in Shikoku during the early 1900s (Ando 2008). The *δ*^34^S values increased as near extinction for otters which were collected in areas with high freshwater accessibility (Figure 3) despite the narrow range of the values (17-19‰). Furthermore, the estimated value indicates high dependence on marine-derived resources for otters though their vibrissae and guard hair would reflect their diet across multiple seasons (Newsome et al. 2009, Naito et al. 2015). Given that Eurasian otters generally change their foraging habitats between freshwater and marine in response to seasonal fluctuation of prey availability or water temperature (Clavero et al. 2006, Kruuk 2006), they had possibly changed in their foraging habitats Japan where is a typical seasonally environment. Thus, our results suggest that otters could not utilize freshwater habitats throughout year.

River degradation also drove otters to extinction through not only prey depletion, but also poisoning due to the accumulation of pesticides, or anthropogenic mortality by local fishery. Top predators including otters are vulnerable to bioaccumulation of toxic substances (Murk et al. 1998, Kruuk 2006). Water pollution due to intensive use of pesticides may have negative effects on otters through bioaccumulation. Further investigation on pesticide concentration in otter hair and vibrissae is needed to understand the negative impacts through bioaccumulation (Iatrou et al. 2019). The most frequent mortality factor of the otter specimens is entanglement by fishery nets, suggesting marine habitats as ecological traps for otters (Barrett et al. 2019). In the study period, coastline habitats might have more abundant prey, but higher human-caused mortality than freshwater habitats. Otters may also prey on fishes within fishery nets easier than those outside the nets because of higher fish density and limiting fish movement.

The current study highlights that freshwater habitat degradation would drive the recent extinction of the last otter population in Japan. Japanese otters initially declined due to commercial fur hunting but never recovered after protection from hunting. Habitat conservation were not option for their protection in Japan at the time. If conservation practice was sufficiently done based on their foraging ecology and habitat requirements, Japanese otters may not extinct by surviving in the remaining suitable habitats. Moreover, many otters had died by a capturing or breeding project for ex situ conservation due to absence of otter specialists (Ando 2008). As establishment of conservation center with otter specialists was key for otter recovery in Europe (Reuther 1995), the involvement of experts played an important role in otter conservation in Japan.

Our study also provides insights into future conservation of other existent *L. lutra* populations and otter species by highlighting the importance of utilization of multiple habitats (i.e., marine and river) for their population viability. Some extant otter species including *L. lutra, Aonyx cinereus* and *Lutrogale perspicillata* have been often observed to alive in human-dominated landscapes (Aadrean and Usio 2017, Tantipisanuh et al. 2023, Tolrà et al. 2024). In Iran, Eurasian otters, which typically inhabit freshwater habitats, have recently been observed coastline in Iran (Karimi and Badelu 2024), suggesting that freshwater degradation caused by land development has forced them to utilize coastal areas. However, it remains unclear whether utilization of or migration to these alternative habitats are sufficient to sustain their population in comparison with the original habitats. Our results highlight that destruction of one major habitat (i.e., freshwater) led the population to depending exclusively on the other (i.e., marine), ultimately preventing its long-term persistence. Thus, when considering the establishment of protected areas for otters, they should encompass multiple interconnected ecosystems to support long-term energy requirements and ensure population persistence. Future research on habitat suitability and connectivity for otters beyond ecosystem boundary needs for establishment of appropriate protected areas for otters worldwide.

## Acknowledgements

We thank Dr. Shuji Yachimori for providing information on otter specimens in the study site. This study was conducted by the supports of JSPS KAKENHI Grant Number 22KK0172 and Joint Research Grant for the Environmental Isotope Study of Research Institute for Humanity and Nature.

