## Supplemental Table S1-S5 for "Freshwater degradation as a potential extinction factor of the Japanese river otters": Supplement_OtterISO.docx

Table S1 *δ*^34^S values of potential food items of the Japanese otter (*Lutra lutra* nippon)

| Categories | Species name | Source | *δ*^34^S values |
| --- | --- | --- | --- |
| Freshwater Fish | *Plecoglossus altivelis* | Freshwater | -2.73882 |
| Freshwater Fish | *Plecoglossus altivelis* | Freshwater | -3.29755 |
| Freshwater Shell | *Semisulcospira libertina* | Freshwater | -2.93236 |
| Freshwater Shell | *Semisulcospira libertina* | Freshwater | -1.88707 |
| Marine-Crab | *Thalamita crenata* | Marine | 21.51716 |
| Marine-Fish | *Trachurus japonicus* | Marine | 18.40863 |
| Marine-Crab | *Thalamita crenata* | Marine | 21.04809 |
| Marine-Shell | *Mytilus coruscus* | Marine | 19.47032 |

Table S2 Summary table of model selection results of the Generalized Linear Models using 5 km buffers

|  | **Model terms (5 km)** | **Residual Deviance** | **AIC** | **ΔAIC** |
| --- | --- | --- | --- | --- |
|  | sex + year + river5 + year: river5 | 22.678 | 88.74 | 0 |
| year + river5 + sex: river5 | | 24.268 | 88.78 | 0.04 |
|  | sex + year + river5 + sex: river5 + year: river5 | 21.967 | 89.79 | 1.05 |
|  | sex + year + river5 | 25.874 | 90.70 | 1.96 |

sex: otter sexes (male or female)

year: year when otter specimen was captured

river5: total length of river within 5km buffer around the sampling points

Table S3 Summary table of model selection results of the Generalized Linear Models using 10 km buffers

| **Model terms (10 km)** | **Residual Deviance** | **AIC** | **ΔAIC** |
| --- | --- | --- | --- |
| year + river10 + year:river10 | 26.436 | 91.34 | 0 |
| sex + year + river10 | 26.809 | 91.76 | 0.42 |
| sex + year + river10 + year:river10 | 25.691 | 92.48 | 1.15 |
| sex + year + river10 + sex:river10 | 26.415 | 93.32 | 1.98 |

sex: otter sexes (male or female)

year: year when otter specimen was captured

river10: total length of river within 10km buffer around the sampling points

Table S4 Summary table of model selection results of the Generalized Linear Models using 20 km buffers

| **Model terms (20 km)** | **Residual Deviance** | **AIC** | **ΔAIC** |
| --- | --- | --- | --- |
| sex | 27.244 | 88.25 | 0 |
| sex + river20 | 27.210 | 90.21 | 1.96 |
| sex + river20 + sex:river20 | 26.412 | 91.32 | 3.07 |
| sex + year + river20 | 26.943 | 91.91 | 3.66 |
| sex + year + river20 + sex:river20 | 25.956 | 92.79 | 4.54 |
| sex + year + river20 + year:river20 | 26.801 | 93.75 | 5.50 |
| sex + year + river10 + sex:river20 + year:river20 | 25.956 | 94.79 | 6.54 |

sex: otter sexes (male or female)

year: year when otter specimen was captured

river20: total length of river within 20km buffer around the sampling points

Table S5 Results of the best model selected by AIC-based model selection using 5 km buffers

| Predictor | Estimate | SE | t value | Pr(>\|t\|) |
| --- | --- | --- | --- | --- |
| (Intercept) | 148.5 | 101.7 | 1.459 | 0.157 |
| Sex (male) | –0.480 | 0.363 | –1.324 | 0.198 |
| Captured year | –0.0668 | 0.0517 | –1.291 | 0.209 |
| **River length** | **–0.00700** | **0.00373** | **–1.874** | **0.0727** |
| **Captured year: River length** | **3.57×10⁻⁶** | **1.90×10⁻⁶** | **1.877** | **0.0722** |

P-values < 0.1 are shown in bold.
